# Multi-colour DNA-qPAINT reveals how Csk nano-clusters regulate T-cell receptor signalling

**DOI:** 10.1101/857516

**Authors:** Sabrina Simoncelli, Juliette Griffié, David J. Williamson, Jack Bibby, Cara Bray, Rose Zamoyska, Andrew P. Cope, Dylan M. Owen

## Abstract

Phosphorylation of the negative regulatory element of the tyrosine kinase Lck by Csk down-modulates T-cell receptor induced signalling. Being constitutively active, Csk spatial organization is responsible for regulating this signalling interaction. Here, we used stoichiometrically accurate, multiplexed, single-molecule super-resolution microscopy (DNA-qPAINT) to image the nanoscale spatial architecture of Csk and two binding partners implicated in its membrane association – PAG and TRAF3. Combined with a newly developed co-clustering analysis framework, we provide a powerful resource for dissecting signalling pathways regulated by spatio-temporal organisation. We found that Csk forms nanoscale clusters proximal to the plasma membrane that are lost post-stimulation and re-recruited at later time points. Unexpectedly, these clusters do not directly co-localise with PAG at the membrane, but instead provide a ready pool of monomers to down-regulate signalling. By generating CRISPR/Cas9 knock-out T-cells, our data also identify that protein tyrosine phosphatase non-receptor type 22 (PTPN22) is essential for Csk nanocluster re-recruitment and for maintenance of the synaptic PAG population.

## Introduction

Protein clustering at the nanoscale appears integral to determine the fate of T-cell receptor (TCR) early activation.^1^ Since clusters are small, they have been extensively studied by single-molecule localization microscopy (SMLM), such as PALM^2^ (photoactivated localisation microscopy) and STORM^3^ (stochastic optical reconstruction microscopy). Previous studies have shown that following TCR engagement, the number and/or density of cell surface receptors (T-cell receptors^4^, integrins^5^), activating kinases (e.g. Lck^6^, ZAP-70^7^) and adaptor proteins (e.g. LAT^4, 8^) clusters generally increases. However, due to the difficulties in quantifying exact protein copy number in SMLM data, different imaging modalities identify diverse protein cluster configurations, ranging from dimers/trimers to larger ‘islands’ and aggregates, opening questions about data interpretation.^9^

Dissecting the role of nanoscale clustering in TCR signalling, therefore, requires to use super-resolution imaging techniques that allows accurate quantification of protein copy number.^9, 10^ In this regard, DNA-PAINT (a variation of point accumulation for imaging in nanoscale topography)^11^, which is a robust SMLM technique that relies on the binding kinetics between two single-stranded DNAs (‘imager’ and ‘docking’ strands), has recently shown impressive advances in quantifying individual molecular targets on synthetic samples that simulate bio-molecular nano-clusters (qPAINT).^12^ This is possible because unlike other SMLM quantification methods, DNA-qPAINT is independent from the dye’s complex photophysics.^13^ Furthermore, DNA-qPAINT enables multi-colour imaging by sequential washout and exchange of imager strands which is not possible with more traditional SMLM.^14^

Csk is the most potent inhibitor of Lck, the primary tyrosine kinase to propagate signalling in T-cells.^15^ Lck adopts an active conformation through the autophosphorylation of Tyr^394^, whereas phosphorylation of its Tyr^505^ residue by Csk promotes its closed, inactive conformation.^16^ Csk exerts a tonic inhibition of membrane-associated Lck in resting T-cells, which appears to set the threshold for TCR activation.^17, 18^ Therefore, for efficient TCR signalling to occur, Csk must translocate away from the membrane-proximal signalling zone to the cytosol.^19^ However, the Lck activation phase only last for minutes before Csk is re-recruited back to the membrane to down-modulate TCR signalling.^19^ Because Csk is constitutively active in T-cells, its relative spatial localization is believed to be critical for regulating TCR signalling. Being a cytosolic protein that lacks any anchor for membrane association, Csk relies on its interaction with transmembrane proteins to achieve this spatial translocation.^20^ In fact, several reports have shown that PAG (phosphoprotein associated with glycolipid-enriched microdomains) phosphorylation at Tyr^317^ controls Csk association and dissociation from lipid rafts at the plasma membrane.^21^ Conversely, TRAF3 (Tumor necrosis factor Receptor Associated Factor), has recently been suggested to act as a chaperone that promotes Csk translocation to the cytoplasm following TCR engagement.^22^ Whereas a large number of studies underline the importance of Csk translocation from the membrane to the cytosol for TCR activation, the underlying mechanism of Csk regulatory duty is still unclear.

Here, we coupled DNA-qPAINT with state-of-the art Bayesian-based cluster analysis approaches to convert raw single-molecule localization data sets to stoichiometrically accurate multi-channel nanoscale protein distribution maps. We use this approach to dissect the functional implications of Csk nano-clustering and spatial organization. We use DNA-qPAINT to quantitatively characterize the nanoscale architecture of Csk, PAG and TRAF3 in resting T-cells and the T-cell immunological synapse. Our data provides new evidence describing, at the single molecule level, how Csk regulates Lck activity, addressing precisely how PAG and TRAF3 are involved. We find that the dynamic nanoscale organization of Csk is indeed a key regulatory element in TCR-induced signal activation and termination, and that it does not involve the formation of 3-component Csk, PAG and TRAF3 complexes. We also show that the absence of Csk’s most relevant binding partner, protein tyrosine phosphatase non-receptor type 22 (PTPN22),^23^ aberrates Csk nano-architecture with consequences for downstream signalling. This work therefore provides a new technology platform for the precise dissection of signalling pathways and new insights into the interplay of kinases and phosphatases in balancing T-cell activation, with implications for understanding immune responses and autoimmune disease.

## Results

### Super-resolution imaging of Csk, PAG and TRAF3 nano-architecture in T-cells using DNA-qPAINT

To characterize the processes whereby Csk down-regulates T-cell receptor (TCR) signalling, we examined the nanoscale organization of Csk, PAG and TRAF3 at different time points in wild-type (WT) Jurkat T-cells proximal to the plasma membrane using DNA-qPAINT, under total internal reflection excitation (Fig. 1). DNA-qPAINT^11^ employs the transient binding of short fluorescently labeled oligonucleotides (‘imager’ strands) to complementary single-stranded DNA targets (‘docking’ strands) chemically coupled to antibodies (Fig. 1a, Supplementary Fig. 1a). While bound, the imager strand can be imaged and localized with nanometer precision. Through sequential imaging of different imager strand sequences targeting different docking strands and their antibodies, DNA-qPAINT overcomes the limitations of other SMLM techniques for multi-colour imaging. To label the proteins for DNA-PAINT imaging, we used monoclonal antibodies conjugated with unique ssDNA docking strand sequences (Supplementary Method 1). We sequentially imaged with the complementary imager strands, each fluorescently labeled with ATTO655, and reconstructed the single-molecule localization maps of each protein (Fig. 1c-e, top).

**Figure 1.**
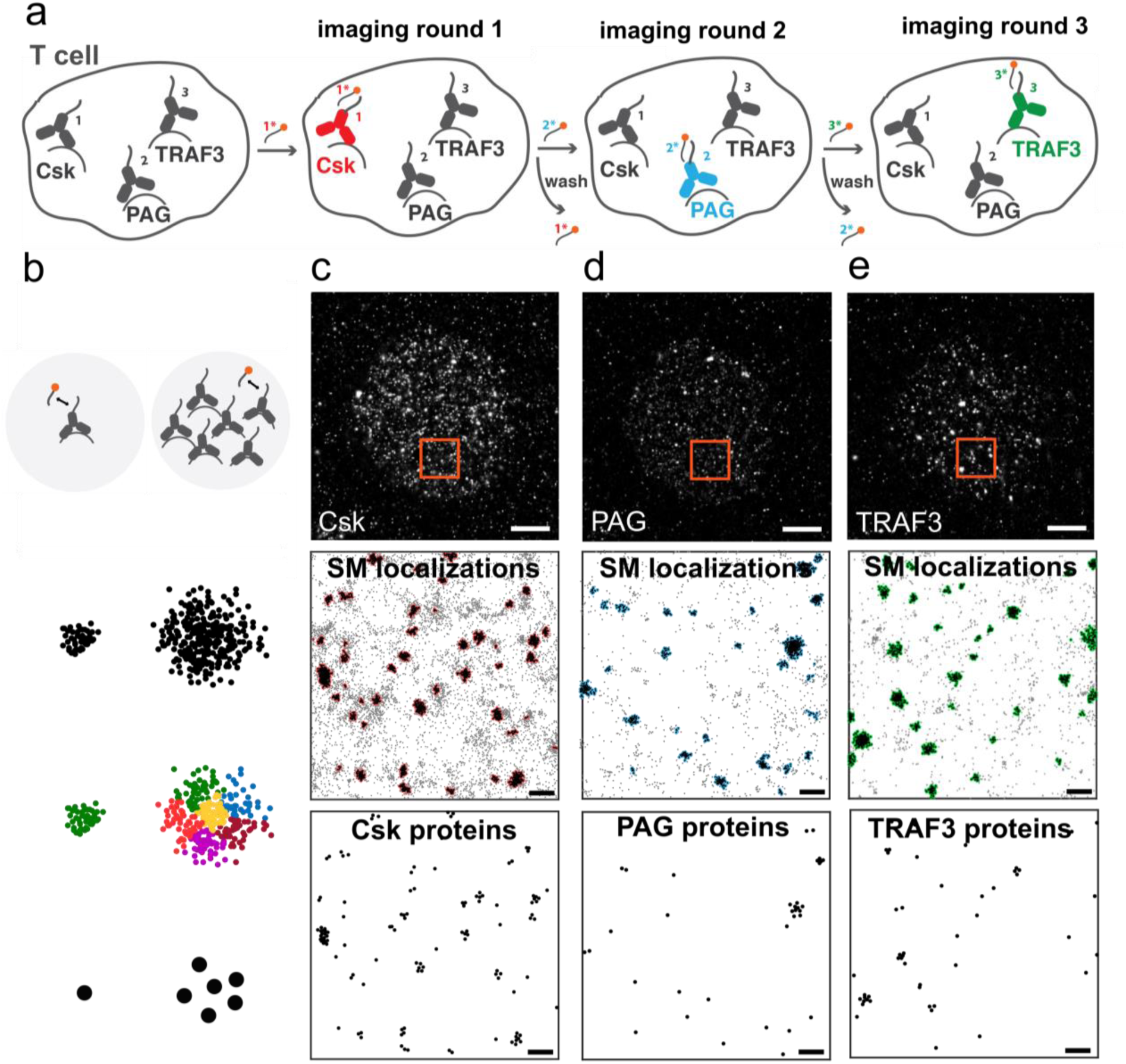
Exchange DNA-PAINT imaging of Csk, PAG and TRAF3 in WT Jurkat T-cells. (**a**) Schematic representation of exchange DNA-qPAINT imaging. Sequential imaging and washing steps are performed until all target proteins are imaged. Super-resolved information is then reconstructed, drift-corrected and aligned to form multiplexed data stacks. (**b**) In DNA-qPAINT, the number of sites can be calculated using the binding kinetic information between fluorescently labeled ‘imager’ strands and their complementary ‘docking’ strands attached to the target antibodies. Using *k*-means function it is then possible to recover the most likely distribution of sites from the single-molecule localizations distribution. (**c**-**e**) DNA-PAINT single-molecule (SM) localization maps (top) of WT Jurkat T-cells supported on poly-L-lysine (PLL) coated glass before they were fixed, permeabilized, and stained with mouse anti-Csk, -PAG and -TRAF3 antibodies each carrying a unique 11 nt single-stranded DNA (docking strand). Images were acquired by sequential imaging with each of the complementary DNA strands labeled with ATTO655 (imager strand) under total-internal reflection illumination. Single-molecule localization distributions (points) were analyzed with a cluster analysis algorithm based on Ripley’s K function, Bayesian statistics and topographical prominence to identify cluster points (red, blue or green outlined) in 4 µm^2^ region maps (center, boxed region). Cluster points were converted into the most likely descriptor of Csk (**c**), PAG (**d**) and TRAF3 (**e**) protein maps (bottom) using the proportionally factor between single-molecule localizations and docking sites and a *k*-means clustering algorithm (Supplementary Figure 1). Scale bars represent 2 µm (top) and 200 nm (center and bottom).

We developed a sophisticated cluster analysis pipeline that allows conversion of DNA-PAINT single-molecule localizations maps (Fig. 1c-e, middle) into absolute protein copy numbers maps (Fig. 1c-e, bottom). In DNA-PAINT, a DNA-coupled antibody is localized a predictable number of times rendering a cluster of localizations around the true position of the protein (Fig. 1b). Using a custom-designed nanoscale DNA-origami platform we calibrated, for each experimental imaging condition, the expected number of localizations per molecular target to be 52 ± 6, and showed that the localization-target relationship is linear over 20 molecular targets with a counting error lower than 10% (Supplementary Fig. 1 and Supplementary Note 1). Non-specific binding events are detected as non-clustered localizations. Thus, after identifying the clusters of single-molecule localizations using a well-established model-based Bayesian cluster analysis method^24, 25^ (colored outlined points in Fig. 1c-e, middle), we used *k*-means clustering to partition the localizations in each detected cluster into their corresponding protein positions (Fig. 1b-e, bottom; Supplementary Fig. 1e). Overall, this approach allows acquisition of a multi-colour stoichiometrically accurate map of protein positions with nanometer spatial precision.

### Nano-clusters of Csk, but monomers of PAG are recruited to the plasma membrane to down-modulate signalling

Using the quantitative approach described above, we reconstructed the distribution of Csk in T-cells on poly-L-lysine (‘control’) and on anti-CD3/anti-CD28 (‘activated’) coated surfaces for either 2 or 8 min (Fig. 2a). To statistically quantify the observed spatial organization, we performed a second stage of Bayesian-based cluster analysis on thirty 2 x 2 µm protein maps regions of interest (ROIs) from ten different Jurkat T-cells (Fig. 2a and 2b). This analysis allowed us to quantify the total number of Csk proteins, the number and size of Csk clusters, and the number of Csk proteins per cluster. The analysis showed that half of the total number of Csk proteins are lost from the plasma membrane-proximal region following TCR engagement (17 ± 3 Csk proteins/µm^2^ compared with 31 ± 5 Csk proteins/µm^2^ in resting cells). At later time points, there is a re-recruitment of Csk back to the membrane, even over-shooting the original, resting protein density (43 ± 7 Csk proteins/µm^2^). The total Csk protein population is composed of two distinct pools: (i) un-clustered Csk proteins, defined as ‘Csk monomers’, or (ii) Csk proteins organized into clusters containing 9 ± 6 Csk proteins per cluster (Supplementary Fig. 2).

Based on this, we therefore assessed the differences between the two pools of Csk proteins (monomers or clusters) during TCR signalling. Immediately after T-cell receptor stimulation, there is a 54% decrease in the number of Csk clusters (0.8 ± 0.5 clusters/µm^2^) with respect to control conditions (1.6 ± 0.9 clusters/µm^2^) (Fib 2b). In late synapses, the number of Csk clusters increases dramatically (3.0 ± 0.7 clusters/µm^2^). Conversely, the number of Csk monomers does not change substantially between resting (15 ± 4 proteins/µm^2^), early (11 ± 2 proteins/µm^2^) and late T-cells synapses (17 ± 4 proteins/µm^2^). Altogether, these data indicate that the early loss of membrane-proximal Csk and its re-recruitment at late times is dominated by the translocation of the clustered Csk pool, rather than the monomeric population.

**Figure 2.**
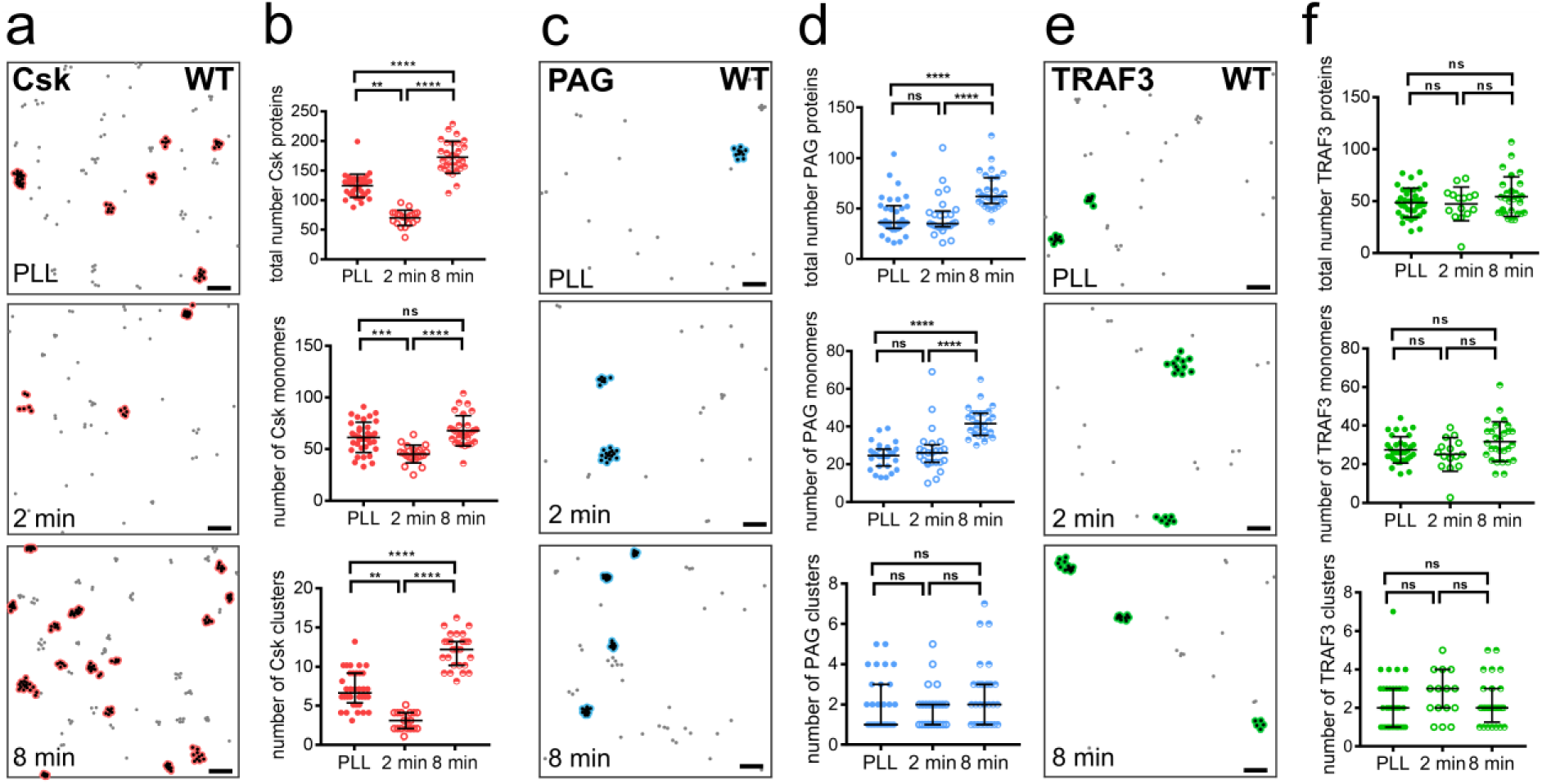
Csk and PAG are recruited back in masse to the membrane during late synapses. Csk (**a**), PAG (**c**) or TRAF3 (**e**) protein maps reconstructed from the localization maps of WT Jurkat T-cells supported on PLL (top) or anti-CD3/anti-CD28 coated glass for either 2 min (center) or 8 min (bottom). For each condition, Csk, PAG or TRAF3 protein maps distributions were analyzed using a Bayesian-based cluster analysis algorithm to identify cluster points (red, blue or green outlined) and extract their properties (proteins per clusters, cluster size, percentage of clustered proteins). Selected descriptors from the cluster analysis of Csk (**b**), PAG (**d**) and TRAF3 (**f**) protein maps representing total number of proteins (top), number of non-clustered proteins (centre) and number of clusters (bottom) per 4 µm^2^ for non-activated (PLL) and activated conditions (2 or 8 min, anti-CD3/anti-CD28). Bars represent means ± SD (**b** and **f**, top & centre) or medians ± interquartile range (**b** and **f**, bottom and **d**). Turkey’s ordinary one-way analysis of variance (ANOVA) (**b** and **f**, top & centre) or Kruskal-Wallis (**b** and **f**, bottom and **d**) multiple comparisons tests, **** P< 0.0001; *** P< 0.0005; ** P< 0.005; ns, not significant. Data are from three independent experiments and represents thirty, 4 µm^2^ regions, obtained from 10-15 cells per condition. Scale bars represent 200 nm.

Since the translocation of Csk is thought to be mediated by PAG and TRAF3, we then characterized the nanoscale organization of these proteins under the same conditions (Fig. 2c-f).^20, 22^ We observed that concomitant with Csk late recruitment, the total number of PAG proteins proximal to the TCR signalosome almost doubles following 8 min of TCR stimulation (from 9 ± 6 to 16 ± 6 PAG proteins/µm^2^) (Fig. 2c and 2d). However, in contrast to the phenomena observed for Csk, PAG is predominantly recruited to the immune synapse as monomers and not as multi-protein assemblies, characterized by a 70% increase in the number of monomers with respect to control conditions. In contrast, clustered PAG was unperturbed post-stimulation, with the number of PAG proteins per cluster and size of PAG clusters remaining unchanged (Supplementary Fig. 3). On the other hand, we found that the overall amount of TRAF3 and its clustering behavior did not significantly change after CD3/CD28 stimulation (Fig. 2e-f and Supplementary Fig. 4).

### Monomeric, not clustered PAG mediates Csk nano-cluster translocation

Many receptors and signalling molecules co-localize into micro- and nano-scale domains during T-cell activation.^1^ We therefore combined multi-colour, stoichiometric super-resolution localization microscopy with Bayesian-based clustering and new co-localization analysis tools to describe the formation of mixed signalling complexes of Csk, PAG and TRAF3 in T-cells.

We first generated three-colour protein maps by assigning a unique pseudo-colour to each species and by aligning and combining each individual colour image using gold nanoparticles as fiducial markers (Fig. 3a, top). Bayesian cluster analysis was performed on the color-coded combined protein maps of thirty ROIs from ten different Jurkat T-cells. Besides retrieving the typical cluster characteristics parameters, we also obtained the information related to the protein composition of each cluster (Fig. 3a, bottom) and plotted it in ternary composition diagrams (Fig. 3b). Every point on the ternary plot represents a different composition of the three components: Csk (red), TRAF3 (green) and PAG (blue). Specifically, we assigned to each cluster a colour code the correlates to the percentage of each protein in the cluster (Fig. 3b) and depicted four circles that correspond to the four possible cluster combinations: (i) Csk-TRAF3; (ii) Csk-PAG; (iii) PAG-TRAF3 and (iv) Csk-TRAF3-PAG. The colour and position of each of these circles relates to the most prominent composition of each cluster combination and the size represents the percentage of that cluster combination in relationship to the others.

**Figure 3.**
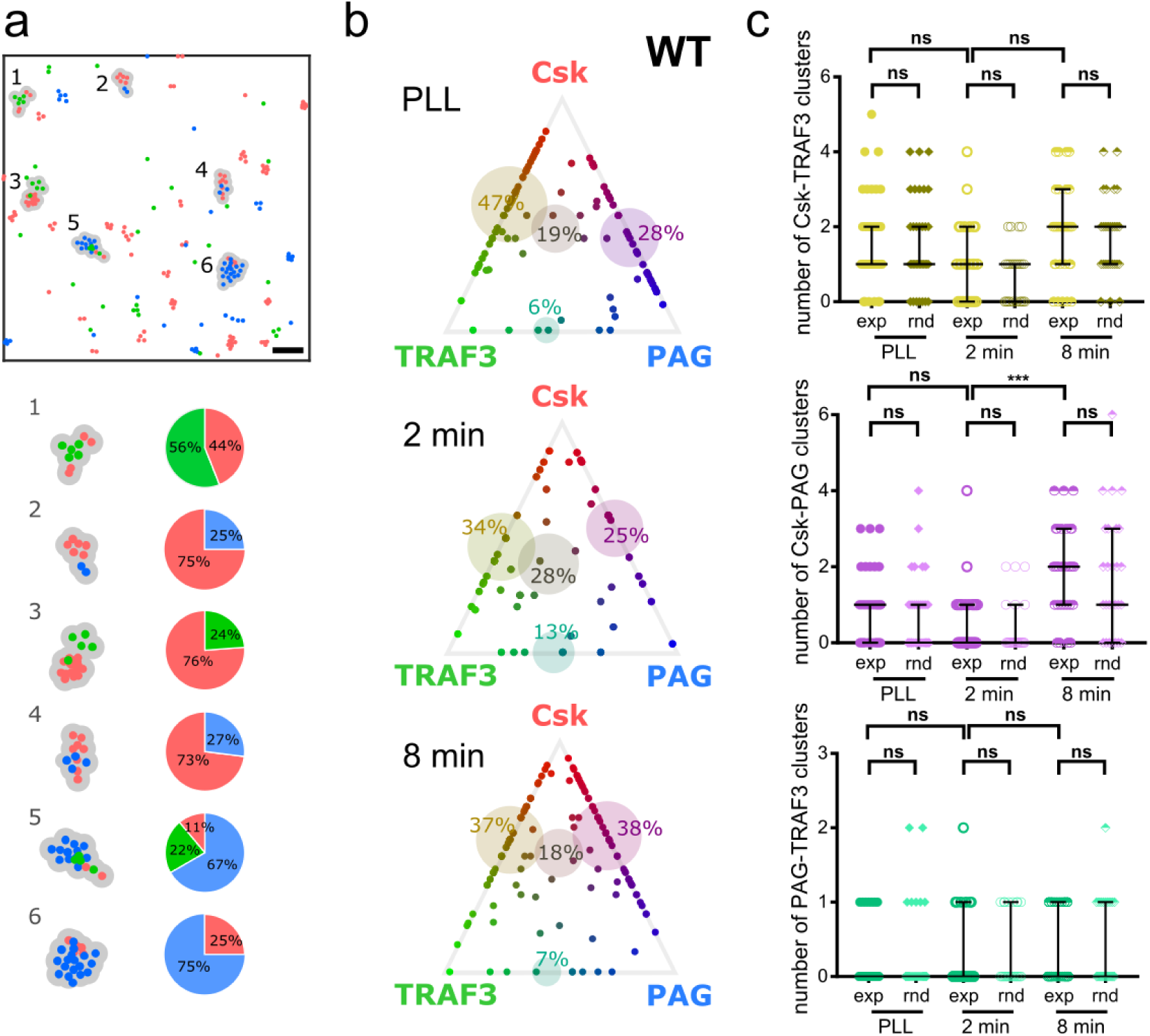
Csk, PAG and TRAF3 mixed protein cluster analysis. (**a**) Exchange DNA-PAINT single-molecule localization maps (top) of WT Jurkat T-cells supported on poly-L-lysine (PLL) coated glass before they were fixed, permeabilized, and stained with mouse anti-Csk (red), -PAG (blue) and -TRAF3 (green) antibodies each carrying a unique 11 nt single-stranded DNAs. Three pseudo-color protein map distributions (points) were analyzed with a Bayesian-based cluster analysis algorithm to identify mixed protein cluster points (grey outlined) in 4 µm^2^ region maps (center, boxed region). The composition of each of the identified cluster was calculated (bottom) and results (**b**) for WT Jurkat T-cells supported on PLL (top) or anti-CD3/anti-CD28 coated glass for either 2 min (center) or 8 min (bottom) were plotted in ternary diagrams. Each point in the ternary plot represents a mixed cluster; the position and color of the point decodes its protein composition. The four main cluster combinations (*i.e* Csk-PAG; Csk-TRAF3; PAG-TRAF3 and Csk-PAG-TRAF3) are represented by a transparent circle positioned in the most likely cluster composition for that combination, and with the size representing the contribution of that cluster combination type with respect to all the found mixed clusters. (**c**) To evaluate the significance of the detected merged clusters, numbers of identified mixed clusters for Csk-TRAF3 (top), Csk-PAG (center) and PAG-TRAF3 (bottom) in experimental data is and TRAF3 at non-stimulated and stimulated conditions (rnd). Bars represent medians ± interquartile range. Kruskal-Wallis multiple comparisons nonparametric test; *** P< 0.0005; * P< 0.05; ns, not significant. Data are from three independent experiments and represents thirty, 4 µm^2^ regions, obtained from 10-15 cells per condition. Scale bar in **a** represents 200 nm.

Ternary graphs for Csk-TRAF3-PAG indicate that both Csk co-clusters with TRAF3 and PAG proteins to a greater extent than any of the other protein combinations (iii and iv), with PAG-TRAF3 co-clusters only contributing to a minority of mixed protein clusters. To quantify the amount of co-clustering which could be the result of coincidental co-localisation, we shuffled the individual thirty ROIs of Csk, PAG and TRAF3 protein maps in each condition and created new merged maps of Csk, PAG and TRAF3 from randomly selected ROIs. We then analyzed the shuffled maps using the same pipeline as described above (Supplementary Fig. 5a). The number of mixed clusters were then counted in each ROI for the different experimental conditions and compared to the corresponding shuffled data (Fig. 3c, exp: experimental vs. rnd: random).

This controlled assessment allowed us to conclude that even the detected Csk-PAG or PAG-TRAF3 cluster combinations do not show any additional, specific co-clustering above that expected for the random case. This indicates that the Csk-PAG interaction does not involve the pool of clustered PAG at the plasma membrane. This is in keeping with our finding that only the monomeric pool of PAG changes following TCR activation. With regards to TRAF3, we did not observe any significant evidence to infer that TRAF3 clusters mediate Csk nano-cluster translocation from the membrane to the cytosol, as there was no significant TRAF3-Csk co-clustering detected.

### PTPN22 is required for the re-recruit Csk to late synapses

Csk is known to be one of the main interaction partners of PTPN22, a key down-regulator of TCR signalling that can dephosphorylate Lck on its active residue, Tyr^394^.^26^ PTPN22 and Csk form a complex that is thought to enable synergistic inhibition of TCR signalling.^26–28^ Other reports, however, suggest that the PTPN22-Csk complex limits PTPN22 function by either restraining its localization to the cytoplasm^29^ or promoting the phosphorylation of an inhibitory residue on PTPN22.^30^ Here, we exploit the high sensitivity of DNA-qPAINT combined with our cluster and co-cluster analysis pipeline to enable, for the first time, a detailed study of how the loss of PTPN22 affects Csk regulatory function.

Recently, we demonstrated the first example of an isogenic human T-cell line that lacks PTPN22 generated using CRISPR gene editing.^31^ We confirmed by western blotting the complete suppression of PTPN22 expression, whilst maintaining constant expression levels of PTPN22 known interactors Lck, Zap-70 and Csk, as well as, PAG and TRAF3 (Fig. 4a).^31^ PTPN22 KO Jurkat T-cells exhibit, in response to TCR stimulation, increased ERK phosphorylation (Fig. 4b) and IL-2 expression levels.^31^ As expected, we do not detect phosphorylation differences in upstream signalling molecules such as Lck and ZAP70 between the WT and PTPN22 KO cells (Supplementary Fig. 6).

**Figure 4.**
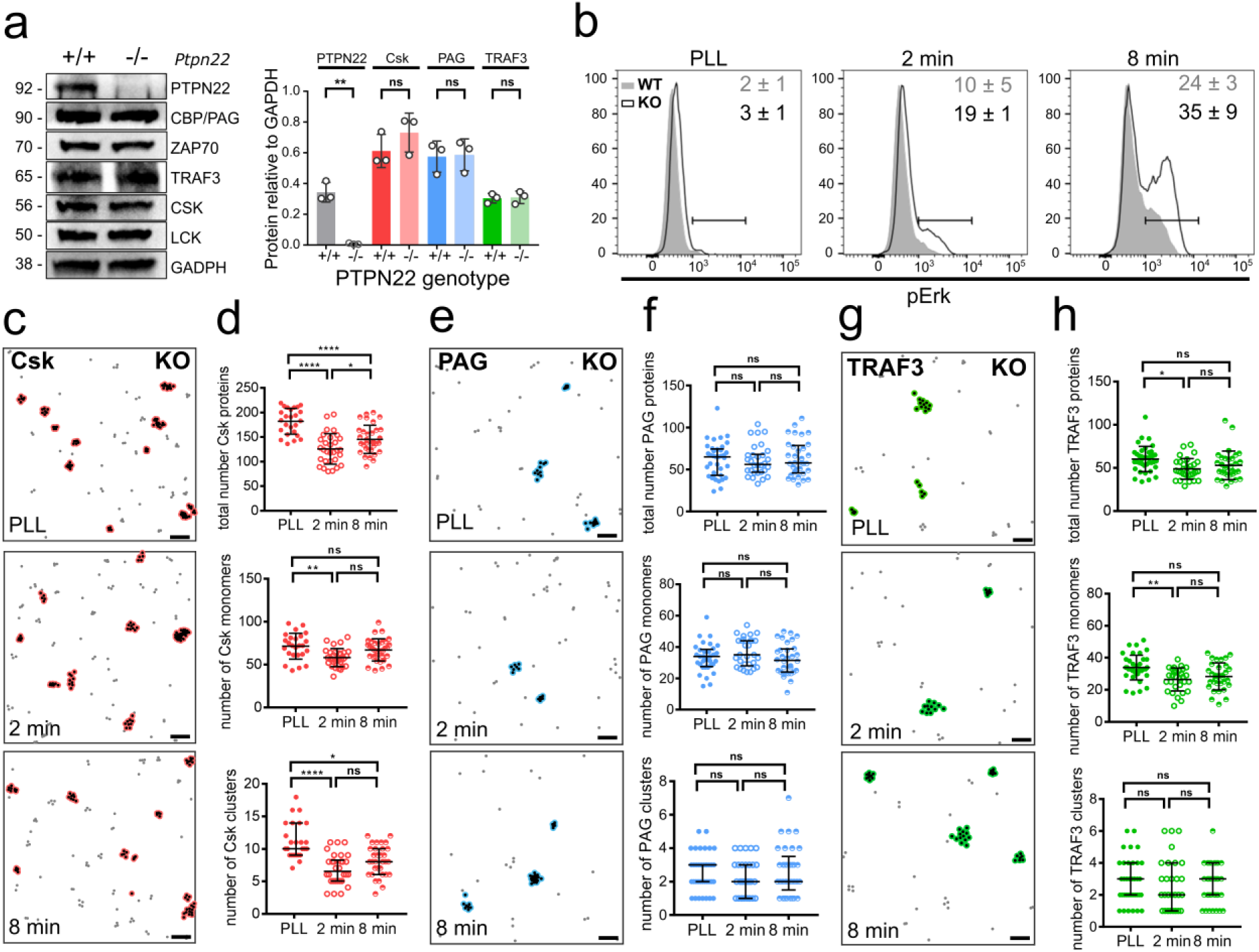
Csk and PAG recruitment at late synapses is impaired in PTPN22 KO T-cells. **(a)** WT and PTPN22 KO T-cells were lysed and analyzed by Western blotting against the indicated proteins. GAPDH was used as loading control. Protein abundance relative to GAPDH in non-stimulated cells was determined. Bars represent means ± SD of three experiments. Turkey’s ordinary one-way analysis of variance (ANOVA): **P = 0.0029; ns, not significant. (**b**) WT (grey) and PTPN22 KO (white) T-cells were supported on PLL or anti-CD3/anti-CD28 coated glass for either 2 min or 8 min before undergoing flow cytometric analysis of Erk phosphorylation. The values represent the percentage of cells that are pErk positive. Data are representative of 3 independent experiments. Csk (**c**), PAG (**e**) and TRAF3 (**g**) protein maps reconstructed from the DNA-PAINT localization maps of PTPN22 KO Jurkat T-cells supported on PLL (top) or anti-CD3/anti-CD28 coated glass for either 2 min (center) or 8 min (bottom). DNA-PAINT imaging and protein maps reconstruction was performed using the same pipeline as described in caption Figure 1. For each condition, protein maps distributions were analyzed using a Bayesian-based cluster analysis algorithm to identify cluster points (red or blue outlined) and extract their properties (proteins per clusters, cluster size, percentage of clustered proteins). Scale bars represent 200 nm. Selected descriptors from the cluster analysis of Csk (**d**), PAG (**f**) and TRAF3 (**h**) protein maps representing total number of proteins (top), number of non-clustered proteins (center) and number of clusters (bottom) per 4 µm^2^ for non-activated (PLL) and activated conditions (2 or 8 min, anti-CD3/anti-CD28). Bars represent means ± SD (**d** and **h**, top & centre) or medians ± interquartile range (**d** and **h**, bottom and **f**). Turkey’s ordinary one-way analysis of variance (ANOVA) (**d** and **h**, top & centre) or Kruskal-Wallis (**d** and **h**, bottom and **f**) multiple comparisons tests, **** P< 0.0001; ** P< 0.005; * P< 0.05; ns, not significant. Data are from three independent experiments and represents thirty, 4 µm^2^ regions, obtained from 10-15 cells per condition.

In contrast to whole-cell level results (Fig. 4a), using DNA-qPAINT, we observed a 46% increase in the total amount of Csk at the membrane with respect to wild-type (WT) cells (Fig. 4c and 4d). This increase is mainly dictated by the higher number of Csk clusters in PTPN22 KO cells (2.5 ± 1.0 Csk clusters/µm^2^) with respect to WT cells (1.6 ± 0.9 Csk clusters/µm^2^), with almost no increase from the monomeric pool of Csk (15 ± 4 Csk proteins/µm^2^ in WT vs. 18 ± 4 Csk proteins/µm^2^ in PTPN22 KO) (*P* values for WT vs KO comparisons are <0.0001 and 0.0410, respectively). As was the case for the WT cells, following TCR stimulation, these Csk clusters are lost, however, the reduction is somewhat less when compared to WT cells (54% reduction in WT vs. 35% in PTPN22 KO). Most interestingly, the absence of PTPN22 leads to a marked failure to recruit Csk clusters back to the plasma membrane during late synapses. In fact, in the KO condition, the number of Csk clusters at late synapses is 20% lower (2.0 ± 1.0 Csk clusters/µm^2^) even than in resting cells (2.5 ± 1.0 Csk clusters/µm^2^). Furthermore, there are no significant differences (*P*-value > 0.9999) between the number of clusters detected at early and late synapses; a high contrast to the behavior observed in the WT cells where the number of Csk clusters quadrupled between early and late activation (*P*-value < 0.0001). Finally, it is worth noting that the clusters characteristics of Csk – *i.e.* number of proteins per cluster and cluster size – in PTPN22 KO cells are very similar to those in WT cells (Supplementary Fig. 2). Taken together, this data indicates that PTPN22 regulates the relative levels of Csk clusters proximal to the plasma membrane by controlling the amount of Csk clusters that migrate there from the cytosol during late synapses.

### PAG late recruitment is also disrupted in the absence of PTPN22

We also examined the effect of PTPN22 deficiency on the nanoscale organization of PAG and TRAF3. Like Csk, PAG in PTPN22 KO cells, showed a marked different behavior in comparison to WT cells. PAG density and clustering before and after TCR stimulation showed no significant differences in the absence of PTPN22 (Fig. 4e and 4f). However, it did display higher basal levels (15 ± 7 PAG proteins/µm^2^) throughout the entire time-course used in this study, even though PAG expression levels at the whole cell level were equivalent for both WT and PTPN22 KO cells (Fig. 4a). With respect to TRAF3 (Fig. 4g and 4h), total TRAF3 at the plasma membrane is higher in PTPN22 KO cells compared to WT (15 ± 4 vs. 12 ± 4 TRAF3 proteins/µm^2^) (*P*-value = 0.0149). Again, these results contrast with whole cell biochemical experiments (Fig. 4a) were we did not observe any difference for TRAF3 expression level between WT and PTPN22 KO cells. Absence of PTPN22 therefore causes a failure to re-recruit Csk at late time points, failure to recruit PAG and an over-abundance of TRAF3 at the plasma membrane. To assess the existence of mixed Csk, PAG and TRAF3 clusters in PTPN22 KO cells we used the same pipeline as described above. Unlike for WT cells, we found that the detection of Csk and TRAF3 mixed clusters in resting cells is significant (*P*-value = 0.0335), and that they separate immediately after activation (Supplementary Fig. 7).

## Discussion

Translocation of Csk to the plasma membrane by transmembrane adaptor proteins, such as PAG, is a critical step towards Src kinase inactivation of the T-cell signalling pathway through phosphorylation of the Lck inhibitory tyrosine site. In resting T-cells, PAG is constitutively phosphorylated at Tyr^317^ enabling binding of Csk through its SH2 domain.^32^ Upon T-cell stimulation, however, PAG becomes dephosphorylated allowing Csk to translocate to the cytoplasm, and as such it alleviates Csk inhibition on active Lck. Csk late re-recruitment back to the plasma membrane is also associated with PAG re-phosphorylation.^19, 32^ Here we found that Csk exists in clusters proximal to the plasma membrane and that it is those clusters which briefly leave the TCR signalosome following early (2 min) TCR stimulation to later (8 min) return to restore its regulatory function. The translocation of Csk clusters to the cytosol enhances the efficacy of downstream signalling by freeing Lck from Csk’s inhibitory action. The re-recruitment of the cytosolic Csk to the membrane is therefore crucial to its function.^33^ The transmembrane adaptor PAG has been long proposed to mediate membrane recruitment of Csk via its SH2 domain.^32^ Our data shows that whilst no significant mixed Csk-PAG clusters exist, the density of both molecules increases at the immune-synapse area during late signalling, with the number of PAG monomers almost doubling. While Csk clusters are not directly associated with PAG, they instead might provide a readily accessible pool of Csk proteins close to the plasma membrane, available for PAG interaction.

We quantified that Csk clusters contain 9 ± 6 Csk proteins; with cluster radius ranging from 18 to 32 nm. This range of cluster parameters agrees very well with other signalling molecules such as TCR (7–30 TCRs per cluster; 35-70 nm cluster radius)^4^ or CD59 (3-9 CD59 per cluster)^34^ as determined by TEM or single-molecule tracking experiments, respectively. Furthermore, they are in line with computational and systems biology approaches which have revealed that cluster size in the range of ∼ 5–10 molecules allows for higher signal fidelity.^35^

We also investigated the role of PTPN22 in Csk clustering dynamics. PTPN22, a key down-regulator of TCR signalling that can dephosphorylate Lck in its active Tyr^394^ residue, is a phosphatase of interest due to a missense mutation (R620W) that has been linked to more than 16 autoimmune diseases in human, including type I diabetes, rheumatoid arthritis and systemic lupus erythematosus.^23, 36^ The C1858T single-nucleotide polymorphism in PTPN22 disrupts its interaction site with Csk.^37^ However, the functional outcomes associated with PTPN22-Csk impaired association remained unclear since both gain- and loss-of phosphatase function has been reported for the PTPN22-R620W variant in humans.^38–40^ By isolating the effect of PTPN22 from the inherent complications in analyzing human data (*i.e.* genetic and environmental variables of distinct individuals), we found that absence of PTPN22 leads to increased ERK phosphorylation and IL-2 expression, which is in keeping with the loss-of-function results observed for the PTPN22 knock-out (KO)^41^ and the orthologous R619W allelic variant in mouse T-cells.^42^

We detected that in the PTPN22 KO cells there is a marked failure to recruit Csk and PAG back during late synapses as well as failure to lose Csk following early TCR engagement. PTPN22 is therefore required to maintain membrane proximal Csk clusters. Quantitative proteomics experiments as well as co-immunoprecipitation results reveal that the PTPN22-Csk association is constitutive; it decreases following TCR stimulation, reaching a maximum after ∼ 10 min of TCR stimulation.^30, 43^ This might indicate that Csk clustering deregulation in PTPN22 KO cells is linked to the loss of Csk-PTPN22 interaction. For example, Veillette et al.^44^ indicated that PTPN22 and Dok adaptors cooperate with PAG to recruit Csk to the plasma membrane to interact with different Src kinases, so that it is possible that PTPN22 might bring Csk to the tyrosine-phosphorylated ITAMs and ZAP-70. In contrast to the results observed in WT cells, we also detected that lack of PTPN22 leads to a modest formation of mixed Csk and TRAF3 clusters in resting T-cells and that there is a slight increase in TRAF3 total expression levels which events out to WT levels following TCR engagement. It is possible that in the absence of PTPN22, TRAF3, which would be otherwise bound to PTPN22 and Csk enhancing their interaction in the cytoplasm,^22^ are now only interacting with Csk leading to increased co-localisation.

In conclusion, we propose a model (Fig. 5) in which Csk nano-clusters are central to increase the effective avidity to membrane-associated proteins, where Csk can exert its inhibitory function on Src family kinases. In this view, Csk clusters provide both, an instant pool of Csk monomers, and also a quick means to exclude Csk proteins from the TCR signalosome to account for proper activation. By excluding Csk nano-clusters from the membrane proximal area, the activation kinase cascade can be sensitive to changes in the localization of only a few ligands. Our model is consistent with experimental evidence from other groups showing that a modest decrease in Csk levels are sufficient to increase Lck phosphorylation on its activating residue, Tyr^394^.^45^ Consistently, by comparing the results in WT and PTPN22 KO cells, we also identified that even only small changes in the total number of Csk molecules, which would be difficult to detect with current biochemical methods, is enough to provoke increase in functional readouts such as IL2 and Erk phosphorylation levels. Overall, our results also illustrate the power of DNA-qPAINT when combined with the latest statistical cluster and co-cluster analysis frameworks for dissecting complex signalling pathways which are regulated by nanoscale spatio-temporal organisation.

**Figure 5.**
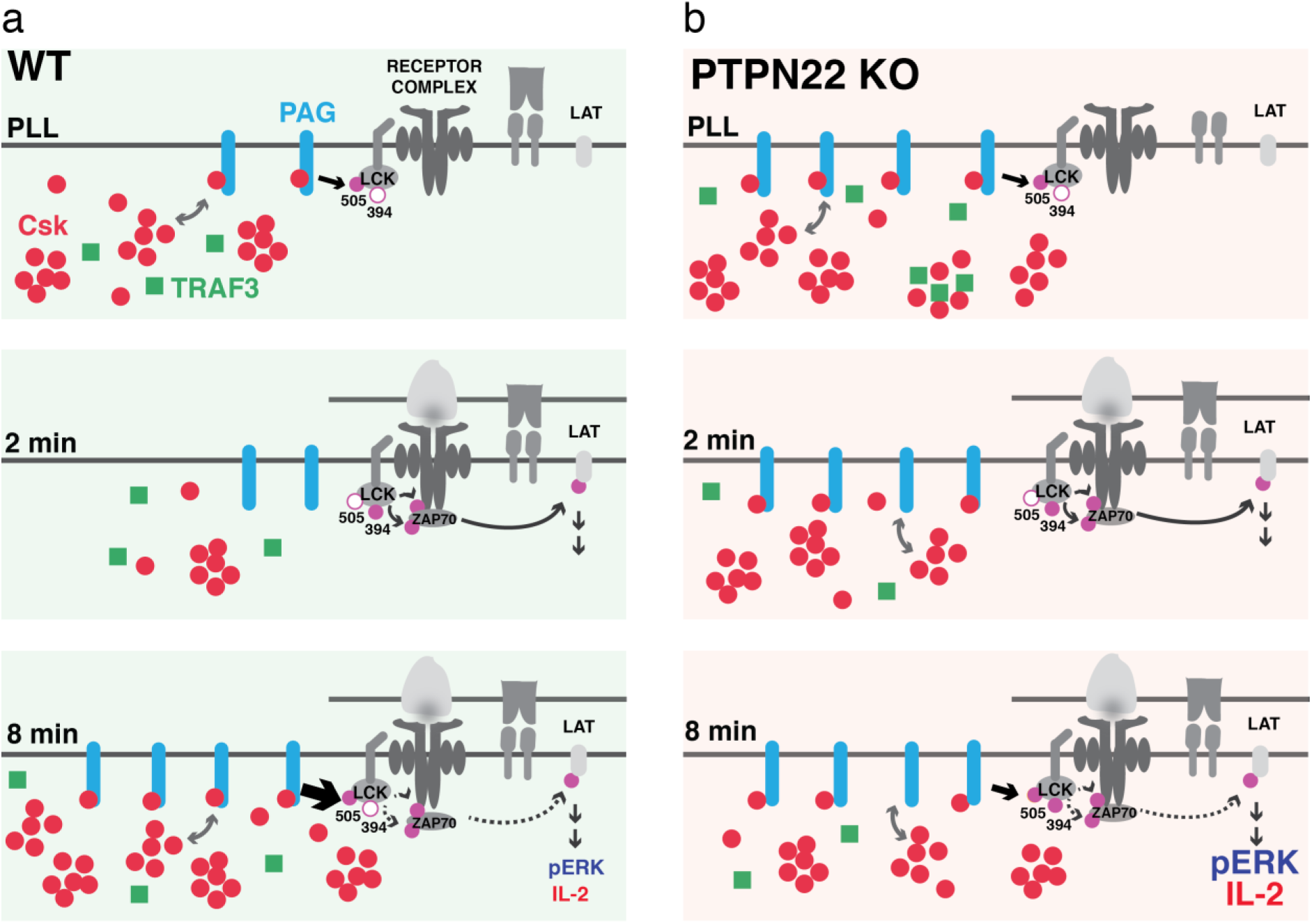
Csk clusters proximal to the plasma membrane deliver Csk monomers to down-regulate signalling in a PTPN22 dependent manner. (a) Csk exerts a tonic inhibitory control of TCR signalling by promoting the closed and inactive conformation of Src-family kinases such as Lck, via phosphorylation of its regulatory sites (top panel). Immediately after T-cell activation, Csk clusters move away from the plasma membrane, which leads to the activation of Lck and initiation of the TCR phosphorylation cascade (middle panel). However, to curtail prolonged T-cell activation, Csk clusters are recruited back *in masse* to the plasma membrane to deliver Csk monomers to late recruited PAG (bottom panel). TRAF3 clusters are, on the other hand, not involved in the translocation of Csk clusters from the plasma membrane to the cytoplasm and vice-versa. (b) Csk and PTPN22 form a complex that modulates the inhibitory function of both proteins in TCR signalling. Loss of PTPN22, leads to impaired Csk clustering dynamics with augmented Csk and PAG expression levels close to the TCR signalosome. Late recruitment of Csk clusters to the plasma membrane is impaired and PAG appears constitutively recruited in PTPN22 KO cells (bottom panel). As a consequence, TCR signalling is augmented and sustained for longer compared to WT T-cells.

## Methods

### Antibodies

Rabbit monoclonal and polyclonal antibodies against PTPN22 (D6D1H) and pLck (Tyr505) were from Cell Signaling Technology and mouse monoclonal antibody against ZAP-70 (29/ZAP70 Kinase) was from BD Biosciences. Mouse monoclonal antibodies against Csk (E-3); Cbp/PAG (G-8); TRAF3 (G-6) were from Santa Cruz, Biotechnology and its conjugation to DNA-PAINT docking strands (Supplementary Table 1) was performed using maleimidePEG2-succinimidyl ester coupling reaction according to a published protocol^11^ as described in Supplementary Method 1. All DNA reagents were from biomers.net GmbH. Secondary anti-rabbit and anti-mouse-HRP antibodies were from Dako.

### Cell lines, media

TCR^−/−^ Jurkat cells were obtained from Hans Stauss, University College London and maintained in Roswell Park Memorial Institute (RPMI-1640, Gibco) medium supplemented with 10% fetal bovine serum (FBS, Serotec) and 5 mM L-glutamine and cultured at 37°C, 5% CO2 in a humidified incubator. Retroviral transduction of TCR constructs was performed as described^46^ and one clone with high expression for a pTax peptide (LLFGYPVYV, derived human T-lymphotropic virus type 1) was selected as the parent line for the Crispr/Cas mediated PTPN22 deletion. PTPN22 KO Jurkat clone was produced by transfecting the selected parent line with plasmids containing Cas9, sgRNA sequences targeting PTPTN22 Exon 1, and a GFP reporter as described in elsewhere.^31^ Western blot and PTPN22 Exon 1 region sequencing confirmed the deletion of PTPN22 protein in the KO cell line. The PTPN22 wild-type (WT) cell line was obtained by transfecting the same parent cell line using only the GFP reporter construct.

### Western blotting

WT and PTPN22 KO T-cells were harvested, washed in PBS and lysed in RIPA buffer (Cell Signaling Technology) with protease inhibitor cocktail (Biolegend). Lysates were added to Laemmli sample buffer (Bio-Rad) with 10% 2-mercaptoethanol, resolved on SDS-PAGE gels and transferred to Immobilon-P polyvinylidene difluoride membranes and blocked with 5% BSA in TBST. After incubation with the corresponding primary and secondary antibodies diluted in 5% BSA in TBST, proteins were visualized by SuperSignal chemiluminescent reaction (Pierce Biotechnology) in a ChemiDoc station (Bio-Rad).

### Phospho flow cytometry

1.5 million WT and PTPN22 KO T-cells per condition were added to a 24-well plate, coated with plate-bound PLL or anti-CD3 (2μg/ml) plus anti-CD28 (5μg/ml). Plates were briefly spun (3 seconds at 300xg) to ensure cells were on the plate bottom, and then placed in the incubator for the indicated time. Cells were fixed with 3% paraformaldehyde, washed, and stained with antigen presenting cells conjugated anti-ERK1/2 pThr202/Thy204 (Biolegend, clone 6B8B69) overnight at 4°C in FOXP3 perm buffer (Biolegend). T-cells were analyzed with a BD LSRFortessa or FACSCanto II (BD Biosciences), and analysis conducted using FlowJo V.10.1 software (Tree Star Inc.).

### Immunofluorescence staining

Six-channel glass-bottomed microscopy chambers (µ-Slide VI 0.5, ibidi) were coated overnight at 4°C with 50 µL of anti-CD3 (8 µg/ml, clone OKT3, Cambridge Bioscience) and anti-CD28 (20 µg/ml, clone CD28.2, BD Bioscience) or for 10 min at room temperature with 0.01% poly-L-lysine (PLL), washed three times in PBS and left in Hank’s balanced salt solution (HBBS) before use. T-cells were resuspended at 3x10^6^ cells/mL in HBBS and immediately added to the coated glass chambers. Following 2 or 8 min of incubation at 37°C, the cells were pH shift-fixed for 90 min with a 4% paraformaldehyde (PFA) and 0.2% glutaraldehyde solution in PEM buffer (80 mM Pipes dipotassium salt, supplemented with 2 mM MgCl2 and 5 mM EGTA, pH 6.8) at 25°C. Following permeabilization for 10 min at 25°C with 0.01% lysolecithin, samples were treated with 20 mM NH4Cl for 10 min at room temperature to quench autofluorescence. The chambers were then block with 5% bovine serum albumin (BSA) for 2 hours and incubated with 0.1 µM DNA-conjugated primary antibodies in 5% BSA in PBS at 4°C overnight. Cells were fixed again with a 2% PFA solution in PBS for 5 min at 25°C, followed by 3x washes with PBS. 100 nm gold nanoparticles (BBI solutions) were added as fiducial markers for drift correction by incubating the sample for 5 min in a 1:2 solution of nanoparticles in PBS and washed 3× washes with PBS, and 3× with PAINT buffer (1x PBS supplemented with 500 mM NaCl). Samples were then used immediately for DNA-PAINT imaging.

### Imaging conditions

Exchange DNA-PAINT imaging was performed on a Nikon N-STORM microscope equipped with a 100× 1.49 numerical aperture oil immersion TIRF objective and a Perfect Focus System. Samples were imaged under TIRF illumination with a 647 nm laser line (∼ 2 kW/cm²) that was coupled into the microscope objective using a quad band set for TIRF (Chroma 89902-ET-405/488/561/647 nm). Fluorescence light was imaged on an EMCCD camera (iXon DU-897U, Andor Technologies) with a final pixel size of 160 nm in the focal plane. For fluid exchange we connected the ibidi µ-Slide VI 0.5 chamber with a silicon tubing (Silicon Tubing 0.5 mm ID, ibidi) via a suitable adapter (Elbow Luer Connector Male, ibidi). Each imaging acquisition step was performed by adding the corresponding 5 nM Atto 655-labelled DNA imager strand in PAINT buffer to the sample (Supplementary Table 1, biomers.net GmbH) followed by a 5 min washing step (corresponding to 10 mL of PAINT buffer). Before the next imager strand solution was introduced, we monitored the camera readout to ensure complete exchange of imager solutions. Sequential imaging and washing steps were performed until all three target proteins (Csk, PAG and TRAF3) were imaged. We acquired, for each imaging step, sequences of 10,000 frames at 10 Hz acquisition rate with an electron multiplier gain of 50 and pre-amp gain profile 1.

### Reconstruction of proteins maps and data analysis

*x,y* localizations coordinates from the raw fluorescent DNA-PAINT imaging experiments were obtained using the ‘Localize’ module of Picasso software.^11^ Localizations with uncertainties greater than 15 nm were removed. Drift correction and multi-colour data alignment was performed using a combination of redundant cross-correlation and fiducial markers approach with the ‘Render’ module of Picasso. Super-resolution image rendering was done by plotting each localization as a Gaussian probably function with standard deviation equal to its localization precision.

To convert the list of *x,y* localizations into a list of *x,y* molecular coordinates we further processed the data using a combination of Bayesian-based cluster analysis algorithms, *k*-means clustering and the prior knowledge of the expected number of single-molecule localizations per docking strand (see Supplementary Note 1). First, we randomly selected thirty non-overlapping 2 µm × 2 µm regions of interest (ROIs) for the analysis of each time-course point experiment (before and following 2- and 8-min TCR stimulation), with a maximum of three ROIs per cell.^24, 25^ To avoid suboptimal clustering results; ROIs were selected such that they do not intersect with cell boundaries. To identify sets of localizations arising from a real docking strand target, we analyzed the data using a robust cluster analysis technique based on Ripley’s K function, Bayesian statistics and topographical prominence. Analysis parameters (prior settings) were choose as default (see Supplementary Table 2) as they genuinely represented the experimental localization-based data sets: (i) cluster-analysis in micron-sized regions; (ii) 20 - 80% of the localizations are in clusters; (iii) radius of the cluster into the 10-500 nm region. Non-cluster points and points attributed to a cluster with fewer than 26 localizations were discarded because they are likely to correspond to noise. This cut-off value was chosen as it is the largest value that gives a cumulative probability of 1 of getting cluster of points with more than that number of localizations (i.e. P (*n* ≥ 26) = 1, where *n* is the number of localizations)). For this consideration we assumed that single molecule localizations are distributed as a Poisson process and we experimentally calibrated that a single docking strand yields, on average, ∼ 52 localizations (see Supplementary Note 1 and Supplementary Fig. 1). To recover the most accurate descriptors (*x,y* molecular coordinates) for the distribution of proteins underlaying the detected distribution of clustered localizations, we implemented a *k*-means clustering strategy in MATLAB (see Supplementary Fig. 1e). The *k*-means algorithm simply partitions a number of observations (i.e. *x,y* localization coordinates per cluster) into *k* sets; returning as a result each set center (i.e. *x,y* molecular coordinates). The value *k*, calculated for each detected cluster of localizations, was estimated to be the ratio between the number of localizations within a cluster and the expected number of localizations per docking strand (see Supplementary Fig. 1c).

To quantify the level and size scale of proteins clustering, we further analyzed the generated *x,y* molecular coordinates with the topographical prominence Ripley’s K based Bayesian model^47^ because it is more robust to detect small multimeric clusters, as it is the case for the observed proteins maps, compared to the originally published Bayesian model script version.^24^ This was possible by choosing priors that increase the typical detectability limit of clusters (around six points per cluster). Where, non-clustered points correspond to monomeric, dimeric or trimeric proteins. Particularly, we defined the priors as having 80% of the points in clusters and we chose a wider radial distribution towards smaller radius values (Supplementary Table 3). This last step it necessary because using *k*-means clustering partitions the data so as to minimize the distances within the sets, and as such, protein cluster sizes will be underestimated.

### Mixed protein cluster analysis

To assess the existence of mixed protein clusters, we merged the stack of generated pseudo-colour molecular localizations coordinates (Csk, PAG and TRAF3) into a single *x,y* coordinate list keeping the identity of the pseudo-colour for each point (merged dataset). Bayesian-based cluster analysis was performed on the merged dataset similar to the one-colour data set as described above; using priors that are bias towards the detection of small mixed protein clusters (Supplementary Table 3). Output data provided both, a label that assigned whether a point was or not in a clustered, with the identity of that cluster, and an extra label identifying the corresponding pseudo-colour of each point. This information permitted allocating to each detected cluster its corresponding protein percentage composition, similar to an RGB code (R = Csk, G = TRAF3 and B = PAG). Mixed protein clusters were considered as such, only when, there were at least two molecular units of each cluster combination type (*i.e.* RG, RB, GB, RGB) in the cluster. Otherwise, the cluster was assigned as pure in the component with higher percentage. To ease graphical representation, each detected mixed protein cluster was depicted as a point in a ternary plot, with the position of the point given by the composition of the three proteins in the cluster. We placed a pure cluster of G (TRAF3) at (*x,y*) = (0,0); of B (PAG) at (1,0) and of Csk at (1/2;√3/2). Resulting in the position of any RGB cluster combination being (*x,y*) = (1/2 * (2B+R)/(R+G+B), √3/2 * R/(R+G+B)). The color of each point in the ternary plot also represents its RGB cluster composition. We depicted a circle for each of the main cluster combinations (*i.e.* RG, RB, GB, RGB) with its position being the most likely cluster composition for that combination, and with the size representing the contribution of that cluster combination type with respect to all the found mixed clusters. To evaluate the significance of the detected merged clusters, we compared them with an artificial merged dataset obtained by randomly mixing the same experimental individual single-colour protein maps and analyzed them using the same pipeline as described above.

### Statistical analysis

Statistical analyses were performed using Prism 6.0 software (GraphPad). The distributions of data points and their variance were determined, and parametric or non-parametric test were utilized as appropriate. Comparison between two groups were evaluated using an unpaired, two-tailed, Mann-Whitney *U* test or Student’s *t* test for normally distributed data. Groups of three or more were compared using the rank-based nonparametric Kruskal-Wallis *H* test or Tukey’s ordinary one-way ANOVA multiple comparison test for normally distributed data. Differences were considered to be statistically significantly different when P < 0.05 for rejecting the null hypothesis.

### Data availability

The data sets generated and analyzed in this study are available from the corresponding author upon reasonable request.

## Acknowledgements

This work has been supported by the BBSRC grant BB/R007365/1 and Versus Arthritis grant 20525. S.S. acknowledges financial support from the Human Frontier Science Program Organization through a postdoctoral fellowship.

## Author contributions

S.S prepared the DNA-coupled primary antibodies, performed the Exchange-PAINT imaging experiments, developed relevant analysis code, analyzed data and wrote the manuscript. J.G. developed clustering and co-clustering analysis code. D.J.W. generated samples. J.B. performed the Western blotting and the flow-cytometry experiments. C.B. generated the Crispr/Cas mediated PTPN22 deletion, while R.Z. supervised the PTPN22 KO Jurkat T-cell construction. D.O. and A.C. conceived the idea and supervised the research. All authors provided feedback on the manuscript.

