## Supplementary material for "Multi-colour DNA-qPAINT reveals how Csk nano-clusters regulate T-cell receptor signalling": Supplemenraty Information and Figures



| | |
|---|---|
| **Supplementary Figure 1** | Calibration with DNA-PAINT imaging. |
| **Supplementary Figure 2** | Analysis of Csk cluster characteristics in WT and PTPN22-KO T cells |
| **Supplementary Figure 3** | Analysis of PAG cluster characteristics in WT and PTPN22-KO T cells. |
| **Supplementary Figure 4** | Analysis of TRAF3 cluster characteristics in WT and PTPN22-KO T cells. |
| **Supplementary Figure 5** | Random merged Csk, PAG and TRAF3 multivalent cluster analysis. |
| **Supplementary Figure 6** | Phosphorylated Lck (Try$^{505}$) and Zap70 (Try$^{319}$) show no significant differences between WT and PTPN22 KO cells. |
| **Supplementary Figure 7** | Csk and TRAF3 coalesce in multivalent clusters in resting PTPN22 KO T cells. |
| **Supplementary Table 1** | DNA-PAINT docking and imager sequences. |
| **Supplementary Table 2** | Priors settings for Bayesian-based cluster analysis of x,y localizations coordinates. |
| **Supplementary Table 3** | Priors settings for Bayesian-based cluster analysis of x,y molecular coordinates. |
| **Supplementary Note 1** | Calibration of DNA-PAINT imaging. |
| **Supplementary Method 1** | DNA-antibody coupling reaction. |
| **Supplementary Method 2** | 2D DNA-origami simulations and analysis. |
| **Supplementary Method 3** | 2D DNA-origami calibration sample and analysis. |

#### Supplementary Figures

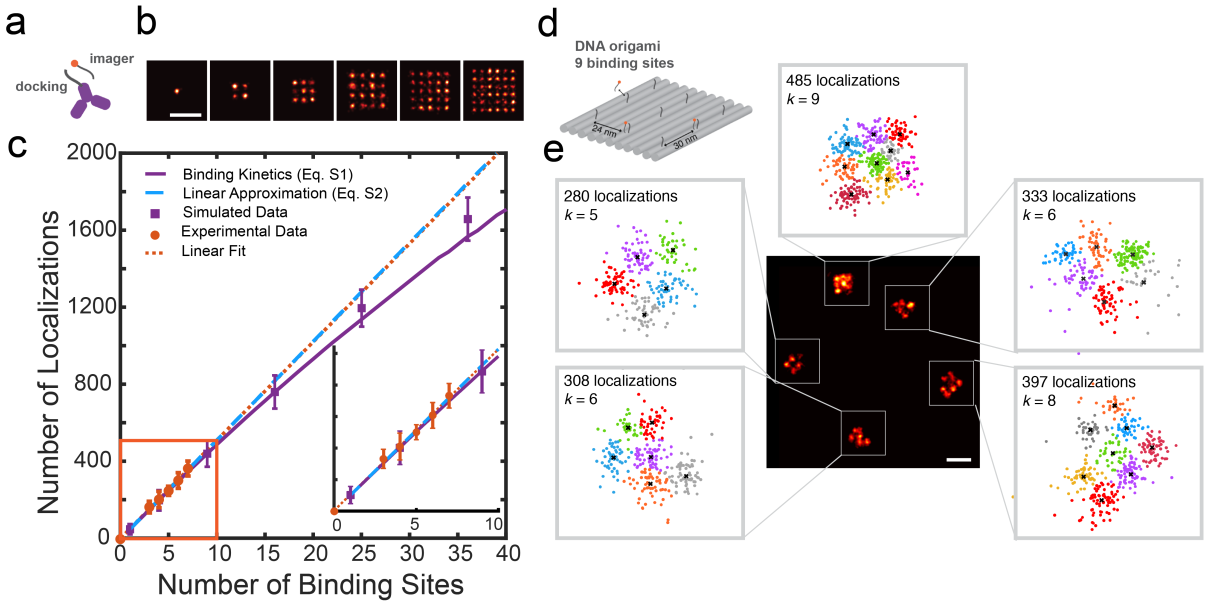


**Supplementary Figure 1. Calibration with DNA-PAINT imaging.** (**a**) Basic concept of DNA-PAINT imaging. Fluorescently labelled 'imager' strands transiently bind to their complementary 'docking' strands attached to a target (antibody, nanobody, etc). Every time an imager strand transiently binds to a target sequence on the sample, it can be imaged and localized. (**b**) Simulated DNA-PAINT images from DNA-origami like structures designed to displayed 1, 4, 9, 16, 25, 36 binding sites spaced 20 nm from each other. Simulations were performed using the 'Simulate' module of Picasso software. Simulation conditions are as specified in Supplementary Method 2. Scale bar represents 50 nm. (**c**) Comparison between in silico simulation (purple squares, mean ± stdev), and in vitro experimental data (orange dots: mean ± stdev; dotted line: linear fit of experimental data, see Supplementary Method 3) for the number of single-molecule localizations detected per DNA origami structure as a function of the number of binding sites observed (visual counting) in each structure. Equation S1 (purple solid line) is in good agreement with both, simulated and experimental data. Zoom-in (orange area) shows deviation from the linear approximation (blue dashed line, Equation S2) already for as little as 10 binding sites, however, the counting error for using the linear approximation is less than 10% for a total of 20 binding sites. (**d**) Schematic of DNA origami structure with 9 designed docking sites (11 nt ssDNA) separated ca. 26 nm from each other. (**e**) DNA-PAINT image (center) and single-molecule (SM) localization maps (magnified views) of the structures (experimental data). Images were acquired using the complementary DNA strand labeled with ATTO655 (imager strand) under total-internal reflection illumination. Using the 'Render' module of Picasso, individual DNA-origami structures were picked, and its number of localizations counted. Single-molecule localization distributions (points) were then partitioned into *k* number of clusters (colored set of points) using the *k*-means clustering algorithm. The *k* value was determined by ratio between the counted number of localizations per origami and the proportionally factor obtained from the linear fit of the experimental data (orange dotted line in **c**): number of localizations = 52 ± 6 x number of binding sites. Scale bar represents 100 nm.

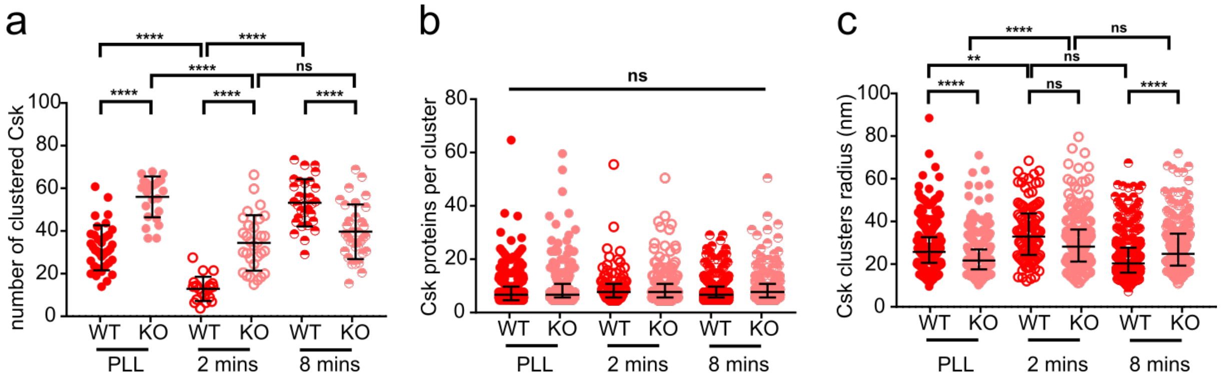


**Supplementary Figure 2. Analysis of Csk cluster characteristics in WT and PTPN22-KO T cells**. Csk protein maps were generated from single molecule localization distributions as described in caption Figure 1 for WT or PTPN22 KO Jurkat T cells supported on PLL or anti-CD3 plus anti-CD28 coated glass for either 2 min or 8 min. Csk protein maps distributions were analyzed using a Bayesian-based cluster analysis algorithm to identify cluster points and extract their properties. Selected descriptors from the cluster analysis of Csk protein maps representing number of clustered Csk proteins (**a**), number of Csk proteins per cluster (**b**) and radius of Csk clusters (**c**) per 4 $\mu m^2$ for non-activated (PLL) and activated conditions (2 or 8 min, anti-CD3 plus anti-CD28). Bars represent means ± SD (**a**) or medians ± interquartile range (**b** & **c**). Turkey's ordinary one-way analysis of variance (ANOVA) (**a**) or Kruskal-Wallis (**b** & **c**) multiple comparisons tests, **** P< 0.0001; ** P< 0.005; ns, not significant. Data are from three independent experiments and represents thirty, 4 $\mu m^2$ regions, obtained from 10-15 cells per condition.

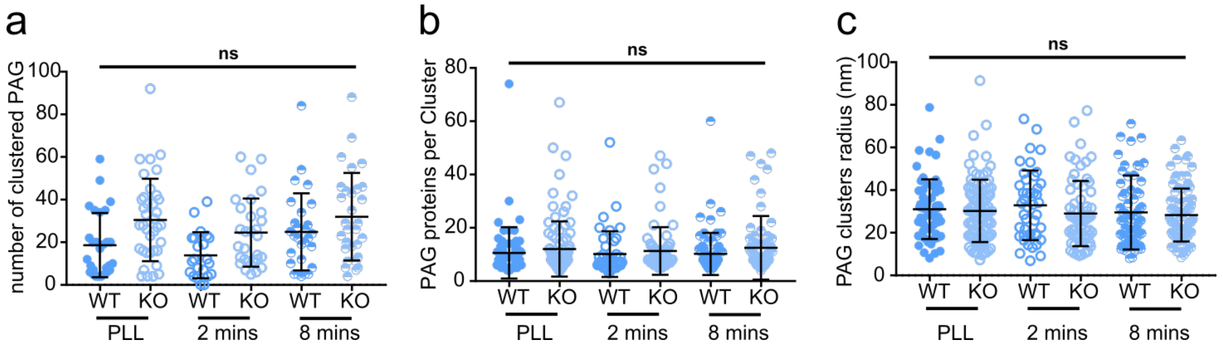


**Supplementary Figure 3. Analysis of PAG cluster characteristics in WT and PTPN22-KO T cells**. PAG protein maps were generated from single molecule localization distributions as described in caption Figure 1 for WT or PTPN22 KO Jurkat T cells supported on PLL or anti-CD3 plus anti-CD28 coated glass for either 2 min or 8 min. PAG protein maps distributions were analyzed using a Bayesian-based cluster analysis algorithm to identify cluster points and extract their properties. Selected descriptors from the cluster analysis of PAG protein maps representing number of clustered PAG proteins (**a**), number of PAG proteins per cluster (**b**) and radius of PAG clusters (**c**) per 4 $\mu m^2$ for non-activated (PLL) and activated conditions (2 or 8 min, anti-CD3 plus anti-CD28). Bars represent medians ± interquartile range. Kruskal-Wallis multiple comparisons nonparametric test; ns, not significant. Data are from three independent experiments and represents thirty, 4 $\mu m^2$ regions, obtained from 10-15 cells per condition.

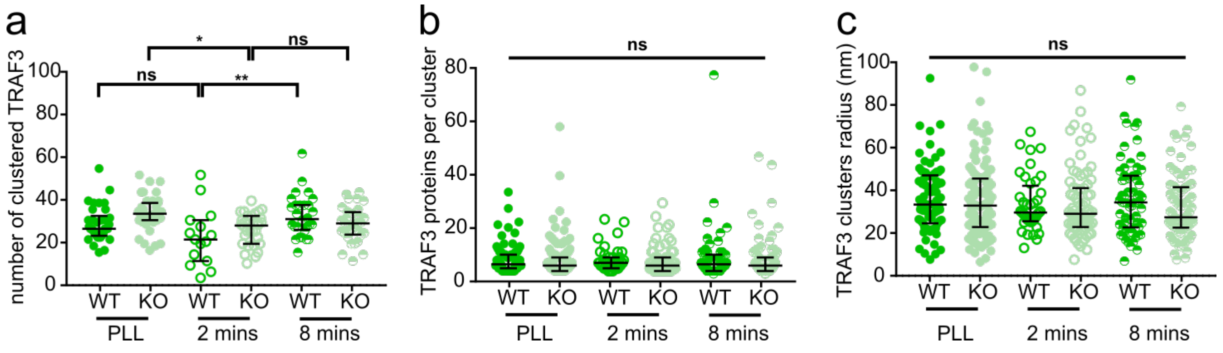


**Supplementary Figure 4. Analysis of TRAF3 cluster characteristics in WT and PTPN22-KO T cells**. TRAF3 protein maps were generated from single molecule localization distributions as described in caption Figure 1 for WT or PTPN22 KO Jurkat T cells supported on PLL or anti-CD3 plus anti-CD28 coated glass for either 2 min or 8 min. TRAF3 protein maps distributions were analyzed using a Bayesian-based cluster analysis algorithm to identify cluster points and extract their properties. Selected descriptors from the cluster analysis of TRAF3 protein maps representing number of clustered TRAF3 proteins (**a**), number of TRAF3 proteins per cluster (**b**) and radius of PAG clusters (**c**) per 4 $\mu m^2$ for non-activated (PLL) and activated conditions (2 or 8 min, anti-CD3 plus anti-CD28). Bars represent means ± SD (**a**) or medians ± interquartile range (**b** & **c**). Turkey's ordinary one-way analysis of variance (ANOVA) (**a**) or Kruskal-Wallis (**b** & *c*) multiple comparisons tests, ** $P < 0.005$; * $P < 0.05$; ns, not significant. Data are from three independent experiments and represents thirty, 4 $\mu m^2$ regions, obtained from 10-15 cells per condition.

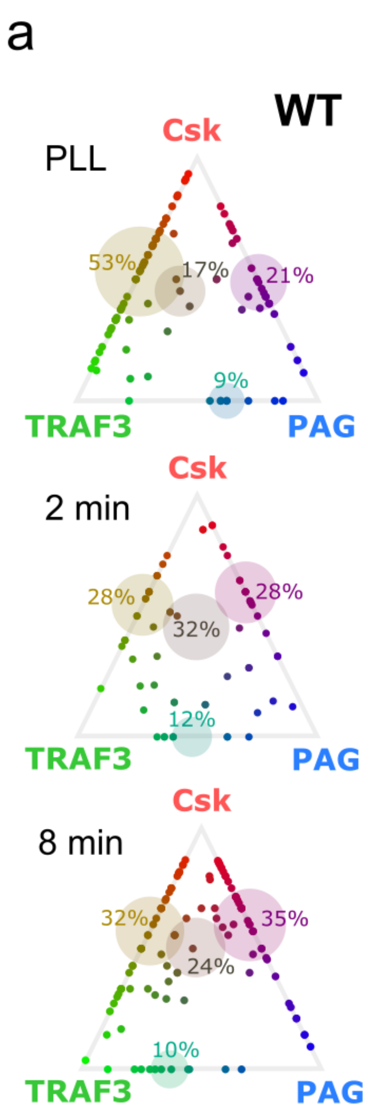


**Supplementary Figure 5. Random merged Csk, PAG and TRAF3 multivalent cluster analysis.** Ternary diagrams for the set of merged data (3-"color" protein map distributions) generated by randomly mixing 4 μm$^2$ protein maps of Csk, PAG and TRAF3 acquired by Exchange DNA-PAINT multicolor imaging in either (**a**) WT or (**b**) PTPTN22 KO Jurkat T cells, at non-stimulated (PLL coated glass, top) and 2 min (center) or 8 min (bottom) stimulated conditions (anti-CD3 + anti-CD28 coated glass). The composition of each detected merged cluster in the shuffle merged data was obtained by analyzing the 3-"color" protein map distributions using the same pipeline as for the experimental set (see caption of Figure 4 and Methods section). Each point in the ternary plot represents a multivalent cluster; the position and color of the point decodes its protein composition. The transparent circles represent the four main cluster combinations (*i.e* Csk-PAG; Csk-TRAF3; PAG-TRAF3 and Csk-PAG-TRAF3) positioned in the most likely cluster composition for that combination, and with the size representing the contribution of that cluster combination type with respect to all the found mixed clusters. Data are from three independent experiments and represents thirty, 4 μm$^2$ regions, obtained from 10-15 cells per condition.

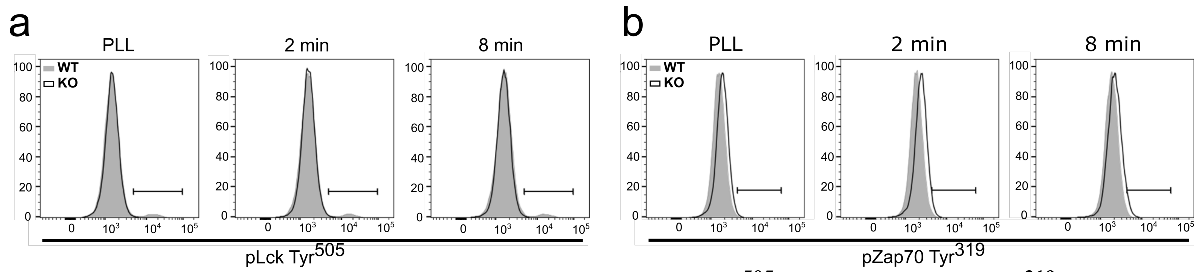


**Supplementary Figure 6. Phosphorylated Lck (Try$^{505}$) and Zap70 (Try$^{319}$) show no significant differences between WT and PTPN22 KO cells**. WT (grey) and PTPN22 KO (white) T cells were supported on PLL or anti-CD3 plus anti-CD28 coated glass for either 2 min or 8 min before undergoing flow cytometric analysis of (**a**) Lck Tyr$^{505}$ and (**b**) ZAP70 Tyr$^{319}$ phosphorylation. Data are representative of 3 independent experiments.

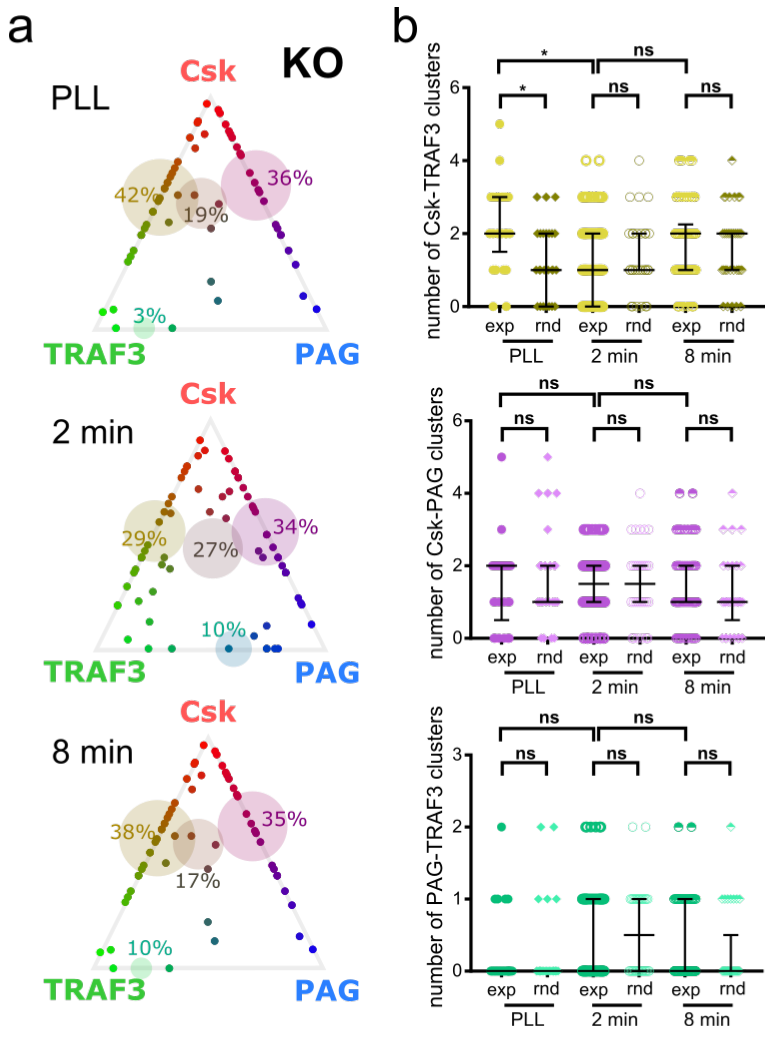


**Supplementary Figure 7. Csk and TRAF3 coalesce in multivalent clusters in resting PTPN22 KO T cells.** (**a**) Ternary diagrams for the set of merged data (3-"color" protein map distributions) of Csk, PAG and TRAF3 acquired by Exchange DNA-PAINT multicolor imaging in either PTPTN22 KO Jurkat T cells, at non-stimulated (PLL coated glass, top) and 2 min (center) or 8 min (bottom) stimulated conditions (anti-CD3 + anti-CD28 coated glass). The composition of each detected cluster was determined using the same pipeline as for the WT data set (see caption of Figure 3 and Methods section). Each point in the ternary plot represents a multivalent cluster; the position and color of the point decodes its protein composition. The transparent circles represent the four main cluster combinations (*i.e* Csk-PAG; Csk-TRAF3; PAG-TRAF3 and Csk-PAG-TRAF3) positioned in the most likely cluster composition for that combination, and with the size representing the contribution of that cluster combination type with respect to all the found mixed clusters. Data are from three independent experiments and represents thirty, 4 $\mu m^2$ regions, obtained from 10-15 cells per condition. (**b**) The significance of the detected merged clusters, numbers of identified multivalent clusters for Csk-TRAF3 (top), Csk-PAG (center) and PAG-TRAF3 (bottom) was calculated by comparing the experimental data set with the results obtained by randomly mixing proteins maps of Csk, PAG and TRAF3 at non-stimulated and stimulated conditions (rnd). Bars represent medians ± interquartile range. Kruskal-Wallis multiple comparisons nonparametric test; * $P< 0.05$; ns, not significant.

#### Supplementary Tables

| Target Protein | Docking Strand | Imager Strand |
|---|---|---|
| Csk | 5'-ATCTAAGTATT-Thiol | ATTO655-TACTTAGATG-3' |
| PAG | 5'-TAGGTAATATT-Thiol | ATTO655-TATTACCTAG-3' |
| TRAF3 | 5'-TATGTAACTTT-Thiol | ATTO655-AGTTACATAC-3' |

**Supplementary Table 1. DNA-PAINT docking and imager sequences.**

| Parameter | Value |
|---|---|
| histbins (nm) | 10,30,50,70,90,110,130,150,170,190,210,230,250,270,290,310,330,350,370,390,410,430,450,470,490,510,530,550,570,590 |
| histvalues | 8,57,104,130,155,168,197,205,216,175,123,91,74,32,24,22,12,11,6,5,3,5,1,3,0,4,0,1,1,1 |
| pbackground | 0.5 |
| alpha | 20 |

**Supplementary Table 2. Priors settings for Bayesian-based cluster analysis of *x,y* localizations coordinates.** The histbins and histvalues priors were chosen to describe a probability distribution function on the cluster radii that best represented the experimental data set. pbackground was chosen as default (0.5) as it has been showed to be robust to true proportions between 0.2 and 0.8. alpha relates to the organization of the clustered points into groups and it was also chosen to be the default value.[1]

| Parameter | Value |
|---|---|
| histbins (nm) | 10,30,50,70,90,110,130,150,170,190,210,230,250,270,290,310,330,350,370,390,410,430,450,470,490,510,530,550,570,590 |
| histvalues | 150,160,180,200,220,220,230,220,216,175,123,91,74,32,24,22,12,11,6,5,3,5,1,3,0,4,0,1,1,1 |
| pbackground | 0.2 |
| alpha | 20 |

**Supplementary Table 3. Priors settings for Bayesian-based cluster analysis of *x,y* molecular coordinates.** The histbins and histvalues priors were chosen to describe a probability distribution function on the cluster radii that best represented the derived molecular coordinated using k-means clustering (*i.e.* descriptors for proteins distribution).

### Supplementary Notes

#### Supplementary Note 1: Calibration of DNA-PAINT imaging

The reversible binding kinetic between docking and imaging strands in DNA-PAINT data allows to predict the number of times that a number of DNA-docking strands (binding sites), $N_{BS}$ , will be visited during the acquisition time ($N_{localizations}$), such that

$$N_{localizations} = \frac{N_{frames} \times N_{BS}}{N_{BS} + \tau_{dark}/\tau_{bright}} \qquad \text{(eq. S1)}$$

where $N_{frames}$ is the number of frames per image stack acquisition; $\tau_{bright}$ is equal to $1/k_{OFF}$ and is in the order of 0.5 – 0.6 s for 9 base-pair double stranded DNA binding; and $\tau_{dark}$ is equal to $1/(k_{ON} \cdot [Imager])$ with $k_{ON}$ being of the order of 2.3 – 1.6 $10^6$ $M^{-1}$ $s^{-1}$.[2] Knowing the experimental concentration of imager strand used and the total number of detected frames; we can estimate the number of localizations that we expect on average for a given number of DNA-docking strands (Supplementary Fig. 1c, purple line). When $\tau_{dark}/\tau_{bright} \gg N_{BS}$, we can use the linear approximation that directly correlates the number of localizations with the number of binding sites as

$$N_{BS} = \frac{N_{localizations} \times \tau_{dark}/\tau_{bright}}{N_{frames}} \qquad \text{(eq. S2)}$$

Equation S2 predicts that for our experimental conditions - *i.e.* a concentration of imager strand of 5 nM and an image stack of 10,000 frames - each docking strand will be detected as a cluster of ca. 40 – 68 localizations depending on exact binding kinetic parameters.

To determine the sensitivity and the dynamic range of the linear approximation, we experimentally imaged (Supplementary Fig. 1e) or stochastically simulated (Supplementary Fig. 1b) single-molecule localization data of DNA origami nanostructures. DNA origami nanostructures offer the possibility to fabricate trillions of identical objects at once while achieving high-specificity and precise positioning of single capturing probes in the nanoscale. As such, they can be used as a breadboard to carry a pre-defined number of DNA-docking strands on their surface (Supplementary Fig. 1b). Experimentally, visual inspection of each DNA origami defines the number of true docking strands incorporated in the structure (binding sites), which can be different for each individual rectangular shape origami. For simulated data, we can input to have 100% incorporation of the docking strands in each of the different type of simulated DNA-origami breadboards (Supplementary Method 2). By comparing the number of docking strands (binding sites) with the number of detected single-molecule localization, we can directly determine how many single-molecule localizations we expect to see per each docking strand. Experimentally, we found that number to be 52 localizations, which correlates with having an association ($k_{ON}$) rate of 1.7 $M^{-1}$ $s^{-1}$ and $\tau_{bright}$ of 0.6 s.

#### Supplementary Methods

##### Supplementary Method 1: DNA-antibody coupling reaction

DNA labelling of a monoclonal primary antibodies was performed using the maleimidePEG2-succinimidyl ester coupling reaction.[3] To reduce the thiolated DNA for the maleimide reaction, 13 µL of the corresponding 1 mM thiol-DNA was incubated with 30 µL of a freshly prepared 250 mM DDT (Thermo Fisher Scientific) solution (1.5 mM EDTA, 0.5x PBS, pH 7.2) on a shaker, in the dark, for 2 h at room temperature. 30 min after the reduction of the thiol-DNA started, 50 µL of 13 µM antibody was incubated with 0.5 µL of 23.5 mM Maleimide-PEG2-succinimidyl ester (Sigma-Aldrich) solution on a shaker, in the dark, for 90 min at 4ºC. Prior DNA-antibody conjugation, both sets of reactions were purified using an Illustra MicroSpin G-25 column (GE Healthcare) to remove excess of DDT and a Zeba desalting column (Thermo Fisher Scientific) to remove excess of cross-linker. Next, both flow-through of the columns were mixed and incubated on a shaker, in the dark, overnight at 4ºC. The next day, DNA excess was removed by Amicon spin filtration (100 kDa). Antibody-DNA concentration was measure with the NanoDrop spectrophotometer and adjusted to 2.5 µM with PBS. DNA-labelled antibodies were store for a maximum of 6 months at 4 °C.

##### Supplementary Method 2: 2D DNA-origami simulations and analysis

For the simulations of Supplementary Figure 1b and c, we used the 'Simulate' module of Picasso software.[4] Simulations were performed using 1, 4, 9, 16, 25 or 36 docking sites on rectangular shape DNA-origami as model input assuming 100% incorporation of the docking strands. Association ($k_{ON}$) and dissociation ($k_{OFF}$) rates were chosen to be 1.7 $10^6$ $M^{-1}$ $s^{-1}$ and 1.7 $s^{-1}$, respectively, as they match with the average number of localizations observed in the our DNA-origami experimental data and they are also in keeping with the typical binding kinetics between 9 base-pair single-stranded DNAs[5]. Imager strand concentration, frame rate and total number of frames were set equal to the experimental conditions: 5 nM, 10 Hz and 10,000 frames, respectively. Simulated data was analyzed using the 'Localize' and 'Render' module of Picasso, which allows quantifying the detected number of localizations per simulated origami. 100 origamis of each type were simulated, and the mean number of localizations was plotted with its corresponding standard deviation as depicted in Supplementary Fig. 1a.

##### Supplementary Method 3: 2D DNA-origami calibration sample and analysis

2D commercial custom made DNA-origami sheets of ~ 70 x 90 nm dimensions (GATTAquant GmbH, Braunschweig, Germany) carrying nine docking (5'-ATTACTTCTTT-3') strands, separated ca. 26 nm from each other, were used to verify the theory and experimentally calibrate the expected number of localizations per molecular target. Biotinylated DNA origami sheets were immobilized on BSA-biotin-neutravidin coated glass-bottomed 6-channel slides (#1.5 glass, µ-Slide VI 0.4, Ibidi) according to the vendor's protocol. Briefly, two buffers were used for the DNA-origami samples immobilization: Buffer A+ comprised of 10 mM Tris-HCl, 100 mM NaCl and 0.05% Tween 20 at pH 8.0 and Buffer B+ which consists of 5 mM Tris-HCl, 10 mM MgCl2, 1 mM EDTA

and 0.05 % Tween 20 at pH 8.0. To coat the surface of the glass slides, first, 80 µl of BSA–biotin solution (1mg/ml in Buffer A+, Sigma-Aldrich) was added to the channel and incubated it for 5 min. After washing the channel with 180 µl of Buffer A+, 40 µl of streptavidin solution (1mg/ml in Buffer A+, Thermo Fisher Scientific) was incubated twice for 5 min. The channel was then washed with 180 µl of Buffer A+ and 180 µl of Buffer B+ prior DNA incubation. DNA-origami stock sample was diluted 1:10 in Buffer B+ and added to the channel for 20 min, followed by 2x washes with 100 µl of Buffer B+. Following DNA-origami immobilization, 100 nm gold nanoparticles fiducial markers (BBI solutions) were added and incubated for 5 min (diluted 1:2 in PBS + 5 mM $MgCl_2$). Gold nanoparticles excess was rinsed 3× with Buffer B+. For TIRF-SMLM imaging, 5 nM fluorescently labeled DNA imager strand (ATTO655-TACTTAGATG-3') was added to the PAINT buffer (1x PBS supplemented with 500 mM NaCl) and loaded into the channel. Samples were then used immediately for DNA-PAINT imaging, using the same experimental setup and conditions described in main Methods section.
*x,y* localizations coordinates and super-resolution images were obtained and reconstructed from the raw fluorescent imaging data as described in the main Methods section. We used the pick tool of the 'Render' module of Picasso to quantify the number of detected localizations per origami and compared it to the number of observable docking sites as showed in Supplementary Figure 1c.
